# mtHsp70 converts mitochondrial proteostasis distress into impaired protein import

**DOI:** 10.1101/2022.09.05.506649

**Authors:** Rupa Banerjee, Vanessa Trauschke, Nils Bertram, Ina Aretz, Christof Osman, Don C. Lamb, Dejana Mokranjac

## Abstract

Functional mitochondria are essential for cell viability and depend on protein import from the cytosol. Impaired protein import initiates various well-characterized cellular programs that rescue or remove dysfunctional mitochondria. However, the molecular mechanism that underlies the initial reduction of protein import into defective mitochondria remained unknown. Here, we found that the redistribution of mtHsp70, mitochondrial chaperone that is involved in both protein import and protein folding, regulates the efficiency of protein import. During early mitochondrial stress, before rescue programs are initiated and membrane potential is affected, mtHsp70-dependent import was specifically impaired and association of mtHsp70 with the import complex reduced. Even under non-stress conditions, the majority of mtHsp70 is found in a substrate-bound state. We propose that the availability of free mtHsp70 limits protein import into mitochondria during stress.

## Introduction

Mitochondria are essential for the overall cellular homeostasis and their function is compromised in ageing and in a number of human diseases (*1-4*). Mitochondria import the vast majority of their proteins from the cytosol (*5*). Recent results show that protein import is also used as a sensitive mechanism to monitor mitochondrial fitness (*3,6,7*). The accumulation of import intermediates in the mitochondrial protein translocases not only prevents import of further proteins into the organelle but also leads to a buildup of mitochondrial precursor proteins in the cytosol, additionally challenging the cytosolic proteostasis network (*8-10*). Over the last years, several cellular pathways were discovered that sense impaired protein import and are activated to counter its potentially fatal consequences. MitoRQC (*11*), mitoTAD (*12*) and mitoCPR (*13*) pathways are activated to clear translocases of stalled import intermediates. Impaired import of PINK1 results in its accumulation on the mitochondrial surface initiating mitophagy, the specific degradation of defective mitochondria by autophagy (*14*). Similarly, the impaired import of ATFS-1 leads to translocation of this transcription factor into the nucleus where it initiates mitochondrial unfolded protein response (mtUPR), a specific transcriptional response with the task of rescuing dysfunctional mitochondria (*15*). More recently, it was discovered that impaired import of DELE1 allows it to bind to and activate the downstream factor HRI, which in turn phosphorylates eIF2α, initiating the integrated stress response (*16, 17*). It is generally accepted that reduced membrane potential is the earliest sign of mitochondrial dysfunction (*3, 6, 7*). The membrane potential across the inner membrane is essential for protein translocation across and into the inner membrane and chemical perturbations that dissipate it, impair protein import and launch various rescue programs. Likewise, a number of mutations that impair mitochondrial function and activate mtUPR also lead to a reduction of membrane potential under steady state conditions (*18*). However, it was reported that mitophagy can be initiated without a detectable change in membrane potential (*19*). It is also unclear how the accumulation of unfolded proteins in the mitochondrial matrix, as a result of either exogenous expression of unfolded proteins or mutations in the components of the mitochondrial proteostasis network, would directly affect the membrane potential across the mitochondrial inner membrane. Thus, the molecular mechanism of how mitochondrial dysfunction is sensed in the first place and converted into a deceleration of protein import remains unclear.

## Results and Discussion

To analyze the early effects of mitochondrial stress, we expressed a mitochondrially-targeted non-folding mutant of mouse dihydrofolate reductase, mDHFR_mut_, in yeast using a regulatable promoter (Fig. 1A). mDHFR_mut_ was equipped with the matrix targeting signal of yeast cytochrome *b*_2_ leading to its localization in the mitochondrial matrix, as described previously (*20*). As expected, long term expression of mDHFR_mut_ impaired the growth of yeast cells (fig. S1A) and led to accumulation of precursor forms of mitochondrial proteins in the cytosol (fig. S1B). At early time points (i.e. 2h after induction), the growth of cells expressing mDHFR_mut_ in the matrix was, however, indistinguishable from that of the control cells (fig. S1A) and no accumulation of precursors was observed (fig. S1B). Also, mitochondrial morphology in control and mDHFR_mut_-expressing cells was virtually indistinguishable at this time point (fig. S1C) indicating that that no obvious harm to the cells has taken place 2h after induction of mDHFR_mut_ expression. To eliminate the confounding effects of other cellular processes and to be able to analyze the effects of the accumulation of unfolded proteins in the matrix directly on mitochondria, we isolated mitochondria from mDHFR_mut_-expressing cells and from control cells 2h after induction of mDHFR_mut_ expression (Fig. 1A). Protein profiles revealed no obvious difference between the two types of isolated mitochondria, except for the presence of mDHFR_mut_ (Fig. 1B). Importantly, we saw no difference in the levels of numerous members of the mitochondrial proteostasis network, including both chaperones and proteases, again suggesting that no rescue program has yet been initiated. We then tested whether the membrane potential of isolated mitochondria was influenced by the presence of mDHFR_mut_ in the matrix. The measured membrane potentials of control and mDHFR_mut_-containing mitochondria were identical (Fig. 1C), confirming the notion that the unfolded proteins in the matrix do not directly affect the membrane potential across the mitochondrial inner membrane. To test whether the presence of unfolded proteins affected the ability of mitochondria to import proteins, we incubated isolated mitochondria with various *in vitro* synthesized, ^35^S-labeled precursor proteins that follow one of the three most frequently used routes into mitochondria *(5)*. Matrix targeted precursors, exemplified here with F1ß and *b*_2_(1-167)ΔDHFR, were imported with lower efficiency in mDHFR_mut_-containing mitochondria as compared to the control ones (Fig. 1D, left panel and fig. S1D, left panel). On the other hand, transport of proteins like *b*_2_(1-167)DHFR and *b*_2_(1-147)DHFR, whose complete translocation into the matrix is prevented due to the presence of a lateral sorting signal that leads to their insertion into the inner membrane, was not affected (Fig. 1D, middle panel and fig. S1D, right panel). Similarly, direct insertion of proteins like the ATP/ADP carrier (AAC) into the inner membrane from the intermembrane space side was unaffected by the presence of unfolded proteins in the matrix (Fig. 1D, right panel). How can these differences in import efficiencies be explained? Import along all three routes depends on the membrane potential *(5)*, which is unaffected in these mitochondria (Fig. 1C), demonstrating that the membrane potential cannot be the cause of the observed differences. The one component of the mitochondrial protein translocation machineries that is specifically required for import into the matrix, the route taken by ca. 50% of all mitochondrial proteins, is mitochondrial Hsp70 (mtHsp70) (*1, 21-23*). As part of the import motor of the TIM23 complex, an estimated 10% of the total mtHsp70 pool mediates translocation of proteins into the matrix at any time (*1, 22, 23*). The majority of mtHsp70 is, however, found in the mitochondrial matrix where it mediates folding of newly imported proteins and prevents their aggregation, as any other member of the Hsp70 family of chaperones. We reasoned that accumulation of unfolded proteins in the matrix could affect import of proteins via mtHsp70.

**Fig. 1.**
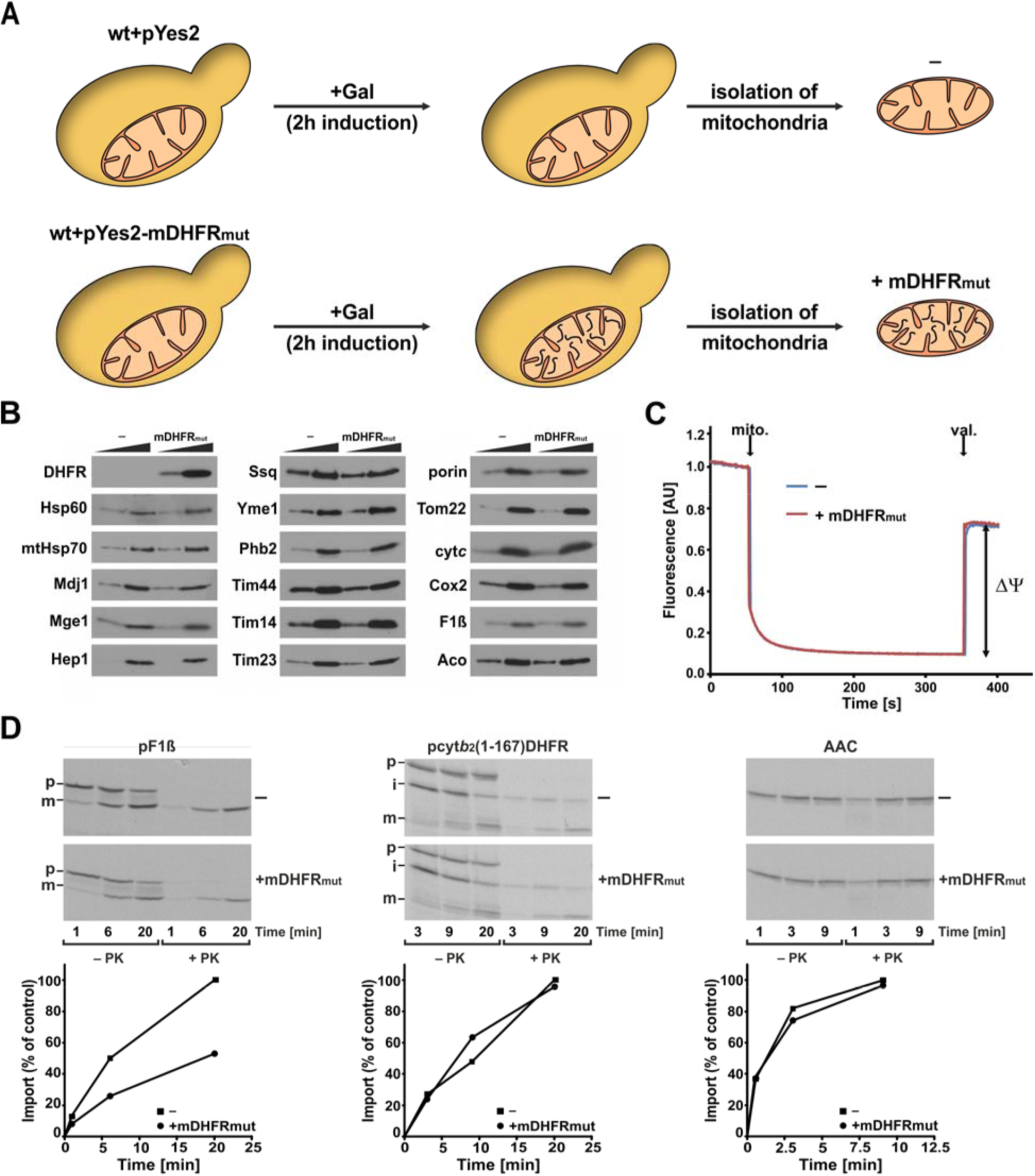
Unfolded proteins in the mitochondrial matrix have no effect on the membrane potential but impair mtHsp70-dependent protein import. (**A**) Schematic overview of the experimental design. Expression of a matrix-targeted non-folding mutant of mouse dihydrofolate reductase, mDHFR_mut_, is induced in yeast cells by addition of galactose. Two hours later, mitochondria were isolated. (**B**) Isolated mitochondria (5 and 20 µg) were analyzed by SDS-PAGE and immunodetection. (**C**) The membrane potential (ΔΨ) of isolated mitochondria (mito.) was measured by fluorescence quenching using DiSC_3_(5). Valinomycin (val.) was added to dissipate the ΔΨ. (**D**) ^35^S-labelled precursor proteins were incubated with isolated mitochondria. At the indicated time points, aliquots were removed and import was stopped. Half of the samples were treated with proteinase K (PK) to degrade the nonimported material. Mitochondria were reisolated and samples analyzed by SDS-PAGE and autoradiography (upper panels). Import reactions were quantified by setting the amount of PK-protected mature protein at the latest time point of import in the control mitochondria to 100% (lower panels). p, precursor; i, intermediate; m, mature forms of mitochondrial proteins.

Hsp70 chaperones consist of an N-terminal nucleotide-binding domain (NBD) and a C-terminal substrate binding domain (SBD) (*24, 25*). The biological functions of Hsp70s are enabled by the conformational changes these proteins undergo as they hydrolyze ATP bound in the NBD and bind substrates in the SBD. Previously, we monitored the conformational cycle of mtHsp70 *in vitro* using single molecule Förster Resonance Energy Transfer (smFRET) spectroscopy by introducing two cysteine residues, one in NBD and the other in SBD, and labeling them with donor and acceptor fluorophores (*26, 27*). This construct is referred to as the “domain sensor”. We used it to show that, in the presence of ATP, the two domains of mtHsp70 are tightly docked onto each other and that, upon ATP-hydrolysis and binding of substrates, the two domains separate ((*26, 27*) and fig. S2A). Building upon this assay, we now developed a method to assess the conformation of mtHsp70 in the physiologically relevant environment of intact mitochondria. To this end, we recombinantly expressed, purified and labeled the precursor form of the domain sensor with its intact mitochondrial targeting signal (Fig. 2A). This fluorescently labelled domain sensor was efficiently imported into isolated mitochondria in a time- and membrane potential-dependent manner, demonstrating that the recombinant protein follows the same pathway into the matrix as the endogenous protein (fig. S2B). After import of the domain sensor, mitochondria remained physiologically active as they were able to generate membrane potential and import further proteins into the matrix (fig. S2C and S2D). To analyze the conformation of mtHsp70 within intact mitochondria, we controlled the concentration of the domain sensor in the import reaction such that, on average, less than one molecule was imported per isolated mitochondrion. The mitochondria were then immobilized on a passivated glass surface and the conformation of mtHsp70 within the organelle assessed on a three-channel single-molecule total internal reflection/widefield fluorescence microscope using highly inclined and laminated optical sheet (HILO) illumination (Fig. 2A, see Materials and Methods for details). Surprisingly, in the fully energized mitochondria, we observed predominantly a low FRET population, indicative of a substrate-bound state, and, to a lesser extent, also a medium FRET state (Fig. 2B). To confirm that the observed low FRET state indeed represents a substrate-bound state of mtHsp70 in intact mitochondria, we performed two additional experiments. First, we analyzed, in intact mitochondria, the conformation of a mutant version of mtHsp70 that is unable to bind substrates. The V459F mutation in mtHsp70 corresponds to the V436F mutation in bacterial Hsp70 DnaK which strongly reduces DnaK’
ss affinity to all substrates analyzed (*28*). The same mutation has no influence on the ATPase activity of mtHsp70 (*29*). When we analyzed the conformation of mtHsp70*V459F* in intact mitochondria, we observed no low FRET peak but rather only the medium FRET population (Fig. 2C). Thus, the mutant that is incapable of binding substrates stably *in vitro* can also not do so in the physiological environment of intact mitochondria. In agreement with this notion, the V459F mutation is lethal *in vivo* (fig. S2E). In a second experiment, we imported fluorescently labelled mtHsp70, either wild-type protein or the V459F mutant, into isolated mitochondria as above, but instead of analyzing their conformation in intact mitochondria, we lysed the organelles with a detergent and analyzed the conformation of the domain sensors in solution. In the presence of ATP, these previously imported mtHsp70s indeed adopted the typical ATP-bound conformation (Fig. 2D). These results show that imported, fluorescently labelled mtHsp70 can adopt the ATP-bound, substrate-free state but it is found predominantly in a substrate-bound conformation in mitochondria, likely due to the presence of ample substrates within the organelle. We conclude that, in physiologically active mitochondria, the majority of mtHsp70 is found in the substrate-bound state, suggesting that the mitochondrial Hsp70 network functions at or near the limits of its capacity. This finding may have broader implications for neurodegenerative diseases as these are frequently caused by a proteostasis collapse within mitochondria.

**Fig. 2.**
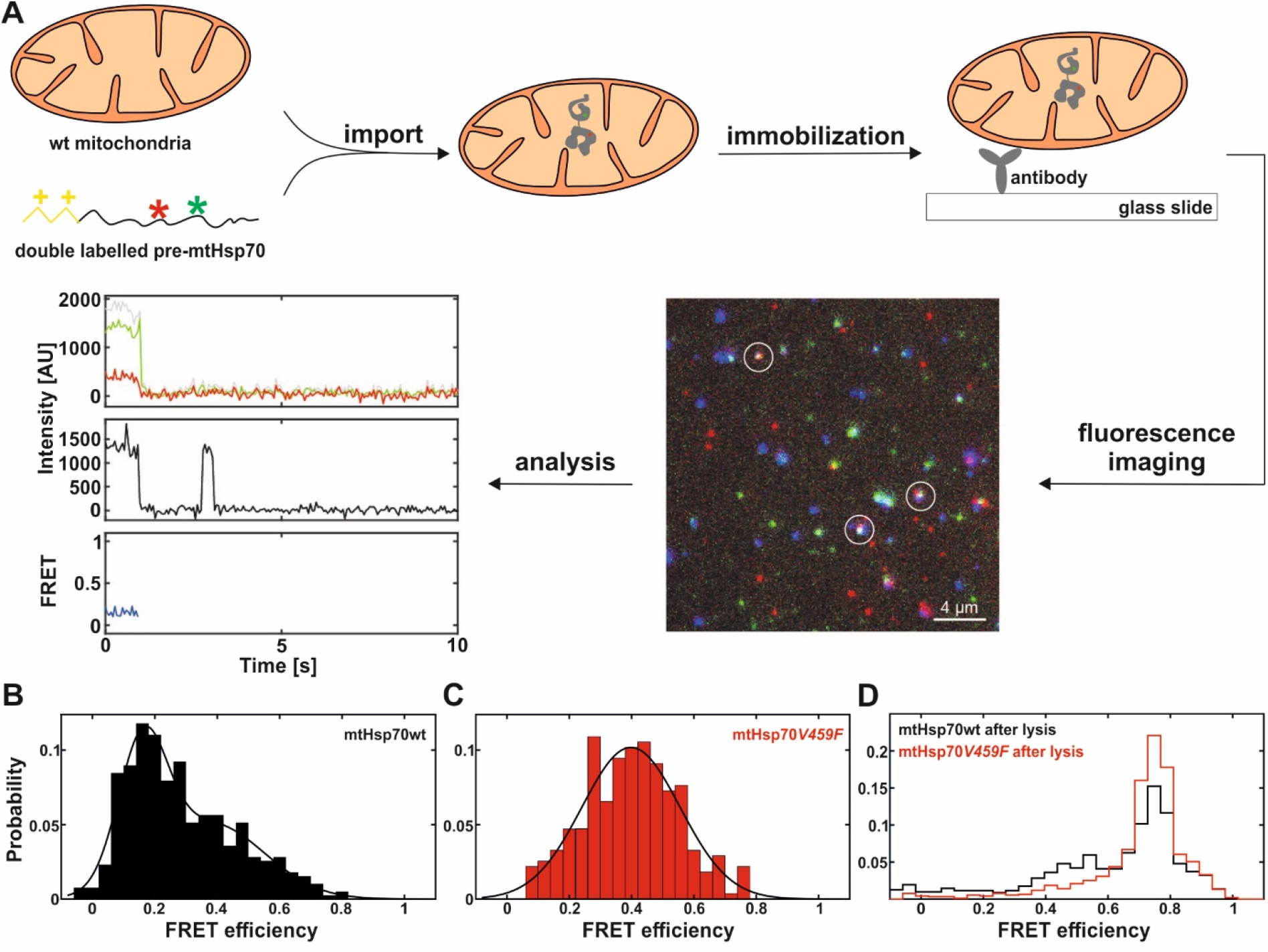
mtHsp70 is predominantly in a substrate-bound state inside mitochondria. (**A**) Schematic overview of the experimental design for single molecule FRET (smFRET) measurements inside isolated mitochondria. A double-labeled mtHsp70, with its matrix targeting signal, was imported into isolated wild-type (wt) mitochondria. Mitochondria were then immobilized on the glass surface and imaged using HILO illumination. The autofluorescence of mitochondria is shown in blue and fluorescence of donor and acceptor fluorophores in green and red, respectively. An overlap of the three signals is visible as white spots and indicates double-labeled protein inside a mitochondrion. From these measurements, single-molecule time traces were extracted. An exemplary single-molecule trace is shown. Top row: ATTO532 (green) and ATTO647 (red) fluorescence intensity after 532 nm excitation. Middle row: ATTO647 fluorescence intensity after 647 nm excitation. Bottom row: calculated FRET efficiency. (**B**) SmFRET histogram of wt mtHsp70 inside energized mitochondria. (**C**) SmFRET histogram of mtHsp70*V459F* inside energized mitochondria. (**D**) Double-labeled wt and *V459F* mutant of mtHsp70s were measured in solution after import, lysis and dilution in buffer containing ATP.

If the majority of mtHsp70 is already present in the substrate-bound state even under non-stress conditions, we reasoned that the presence of additional unfolded proteins in the matrix may decrease the availability of mtHsp70 for the TIM23 complex. To this end, we first immunoprecipitated mtHsp70 and its interaction partners from control and mDHFR_mut_-containing mitochondria. Association of mtHsp70 with the components of the TIM23 complex, exemplified here by Tim17 and Tim14, was reduced in the presence of mDHFR_mut_ (Fig. 3A). The same was observed for the association of mtHsp70 with Hep1, a dedicated chaperone for mtHsp70 (*30*). Concomitantly, mtHsp70 was found to bind to mDHFR_mut_ (Fig. 3A). Next, we analyzed the crosslinking pattern of Tim44, the subunit of the TIM23 complex that recruits mtHsp70 to the translocase (*31*), in intact mitochondria. In energized mitochondria, Tim44 can be crosslinked to mtHsp70 whereas, upon ATP-depletion, the intensity of the Tim44-mtHsp70 crosslinks diminishes (Fig. 3B and (*31, 32*)). In mitochondria containing mDHFR_mut_ in the matrix, the crosslinks of Tim44 to mtHsp70 were dramatically reduced, both in the presence and in the absence of ATP. In contrast, the crosslinks of Tim44 to Tim14 and Tim16, other subunits of the TIM23 complex, were unaffected. Taken together, these results indicate that unfolded proteins in the mitochondrial matrix interfere with the interaction of mtHsp70 with the TIM23 complex, providing a likely explanation for the observed impairment of mtHsp70-dependent protein import into mitochondria.

**Fig. 3.**
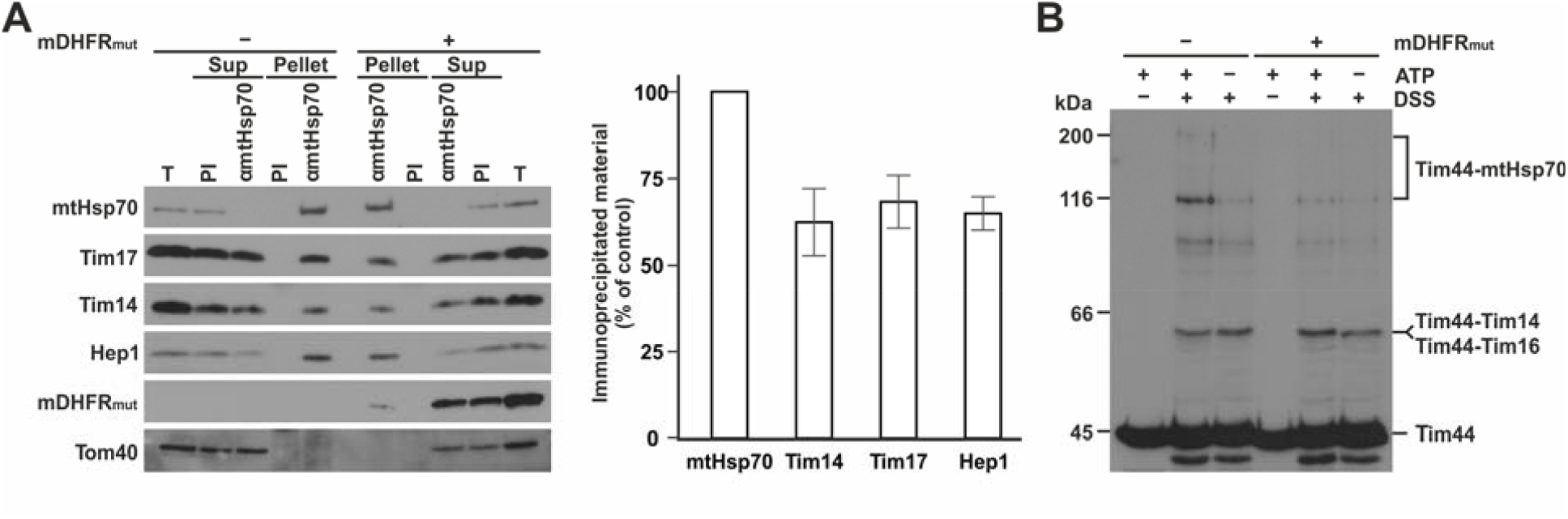
Unfolded proteins in the mitochondrial matrix remove mtHsp70 from the TIM23 complex. (**A**) Immunoprecipitation of mtHsp70 and its interacting proteins from mitochondrial lysates. Antibodies present in a preimmune serum (PI) were used as a control. Total (T, 20%), supernatant (Sup, 20%) and pellet (100%) fractions were analyzed by SDS-PAGE and immunodetection using the indicated antibodies (left panel). Quantification of immunoprecipitated material (right panel). The amount of immunoprecipitated material in control mitochondria was normalized to the amount of immunoprecipitated mtHsp70 and set to 100%. Mean ± s.e.m (n=6). (**B**) Crosslinking in intact mitochondria with disuccinimidyl suberate (DSS). ATP levels of isolated mitochondria were manipulated prior to the addition of the crosslinker. After quenching of excess crosslinker, mitochondria were reisolated and samples analyzed by SDS-PAGE and immunodetection using Tim44 antibodies. Previously identified crosslinks of Tim44 are indicated.

The data obtained here suggest a molecular mechanism of how cells use protein import to sense mitochondrial dysfunction (Fig 4). Under normal conditions, a fraction of mtHsp70 binds to the TIM23 complex and mediates protein import. The majority of mtHsp70 is in the matrix where it is involved in the folding of proteins and prevention of protein aggregation. When the number of unfolded proteins in the matrix increases, some of mtHsp70 will be redistributed from the TIM23 complex to deal with the increased folding-need in the matrix. Removal of mtHsp70 from the TIM23 complex will, in turn, reduce the ability of mitochondria to import further proteins into the matrix. Reduced import efficiency of effector proteins will lead to their accumulation on the mitochondrial surface or elsewhere in the cell allowing them to initiate rescue programs. As the mtHsp70 network functions at the limits of its capacity, even the smallest proteostasis problems in the matrix can be easily detected, allowing the cell to promptly react and prevent potentially fatal consequences of mitochondrial dysfunction.

**Fig. 4.**
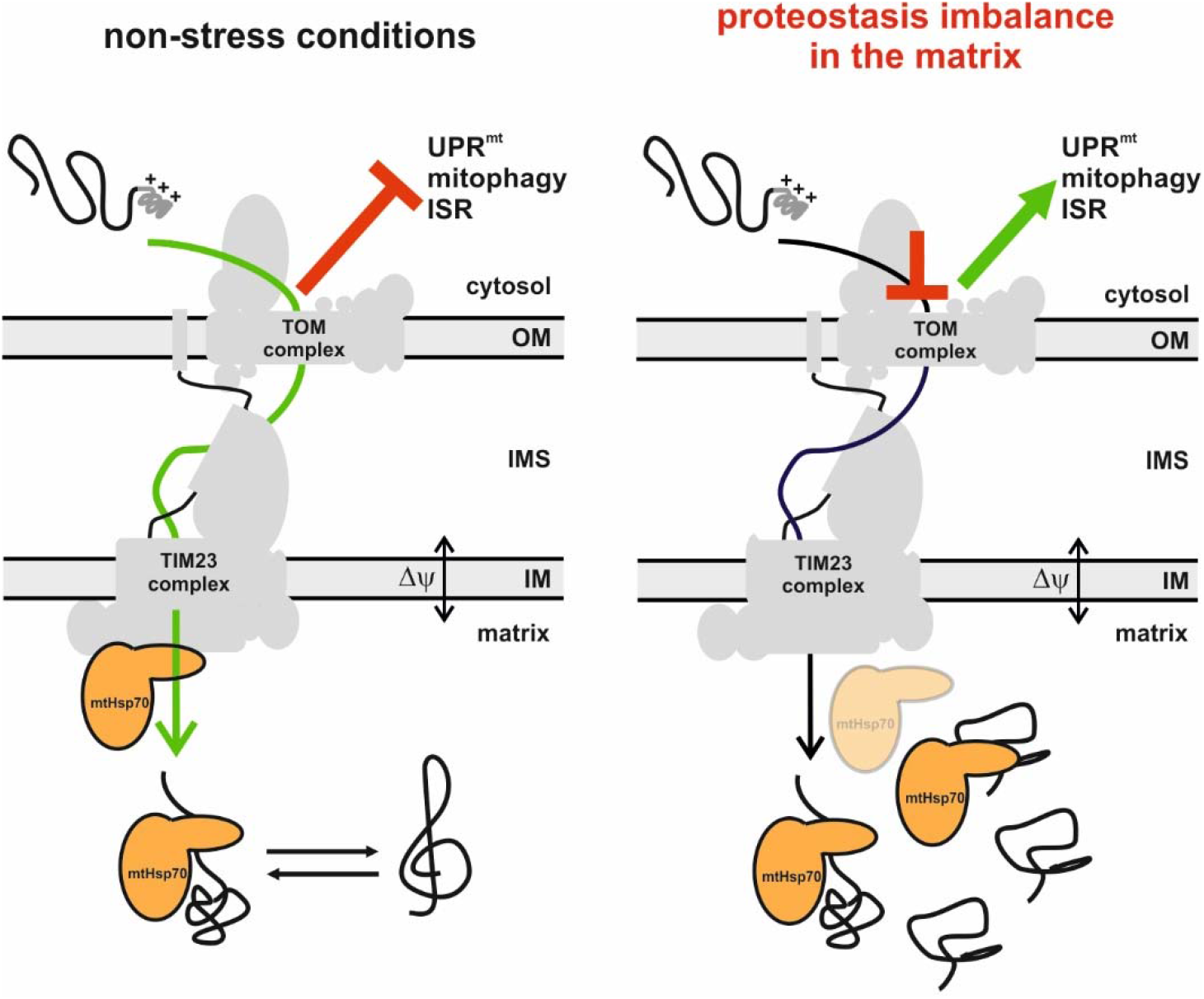
mtHsp70 converts mitochondrial proteostasis distress into impaired protein import. *Left panel:* Under normal conditions, mtHsp70 mediates import of proteins as part of the TIM23 complex and is involved in folding of proteins in the matrix. *Right panel:* Mitochondrial stress, caused by for example increased abundance of unfolded proteins in the matrix, leads to a dissociation of mtHsp70 from the TIM23 complex. This, in turn, decreases protein import. Decreased import of effector proteins initiates rescue programs. In this way, mtHsp70 monitors mitochondrial fitness and converts it into the efficiency of protein import.

## Acknowledgments

We thank Petra Robisch and Zdenka Stanic for the expert technical assistance. We acknowledge the Biophysics Core Facility of the BMC for use of Typhoon FLA9500. This study was supported by the Deutsche Forschungsgemeinschaft (MO1944/1-2, MO1944/2-1 and NE101/28-1 to D.M. and SFB1035, Project A11 to D.C.L.) and German Academic Exchange Service doctoral fellowship to R.B.

## Author contributions

Conceptualization: RB, VT, DCL, DM

Methodology: RB, VT, NB, IA, CO, DCL,DM

Investigation: RB, VT, NB, IA, CO, DCL, DM

Visualization: RB, VT, NB, IA, DM

Funding acquisition: DCL, DM

Project administration: DCL, DM

Supervision: CO, DCL, DM

Writing – original draft: RB, VT, DCL, DM

Writing – review & editing: RB, VT, NB, IA, CO, DCL, DM

---

**Fig. S1.**
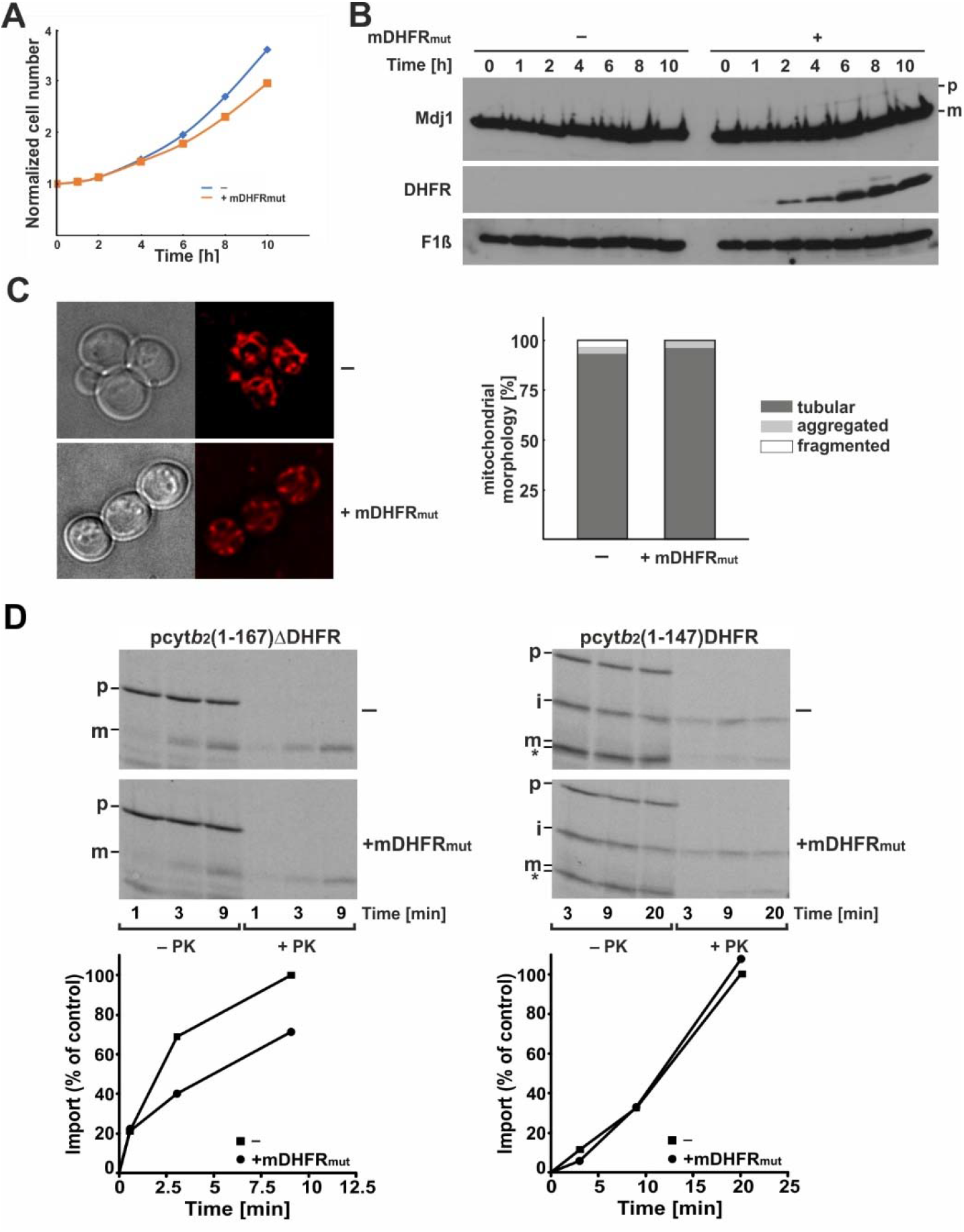
Expression of unfolded proteins in the mitochondrial matrix. (**A**) Prolonged expression of unfolded proteins in the mitochondrial matrix impairs the growth of yeast cells. Yeast cells harboring a plasmid for a regulatable expression of a matrix-targeted unfolded protein, mDFHR_mut_, or the empty plasmid as a control, were grown in a logarithmic growth phase for two days. Galactose was then added to the growth medium to induce the expression of the unfolded proteins and cell growth was monitored with time. (**B**) At the indicated time points after addition of galactose, aliquots were taken from the cultures, whole cell extracts prepared, and analyzed using SDS-PAGE and western blotting using the indicated antibodies. (**C**) Two hours after addition of galactose, MitoTracker was added to the yeast cell cultures to stain mitochondria. Yeast cells were immobilized on agar pads and analyzed using fluorescence microscopy. A typical image is shown in the left panel. A blind quantification of the mitochondrial morphology into tubular, aggregated and fragmented structures was performed (right panel). Ca. 400 cells per strain were analyzed. (**D**) Import of ^35^S labelled precursor proteins into mitochondria isolated from cells two hours after switching to galactose-containing medium. The uptake of two precursor proteins was followed as a function of time: pcyt*b*_2_(1-167)ΔDHFR – a chimeric protein consisting of the first 167 residues, with an inactivated lateral sorting signal from yeast cytochrome *b*_2_, fused to mouse dihydrofolate reductase and pcyt*b*_2_(1-147)DHFR – a chimeric protein consisting of the first 147 residues of yeast cytochrome *b*_2_ fused to mouse dihydrofolate reductase. The former precursor is imported into the matrix in a membrane-potential-dependent and mtHsp70-dependent manner and the latter is laterally inserted into the inner membrane in a membrane potential-dependent but mtHsp70-independent manner. At the indicated time points, aliquots were removed from the import reactions and import stopped. Half of the samples was treated with proteinase K (PK) to degrade the nonimported material. Mitochondria were reisolated and samples analyzed by SDS-PAGE and autoradiography (upper panels). Import reactions were quantified by setting the amount of PK-protected mature protein at the latest time point of import in control mitochondria to 100% (lower panels). p, precursor; i, intermediate; m, mature forms of the mitochondrial proteins.

**Fig. S2.**
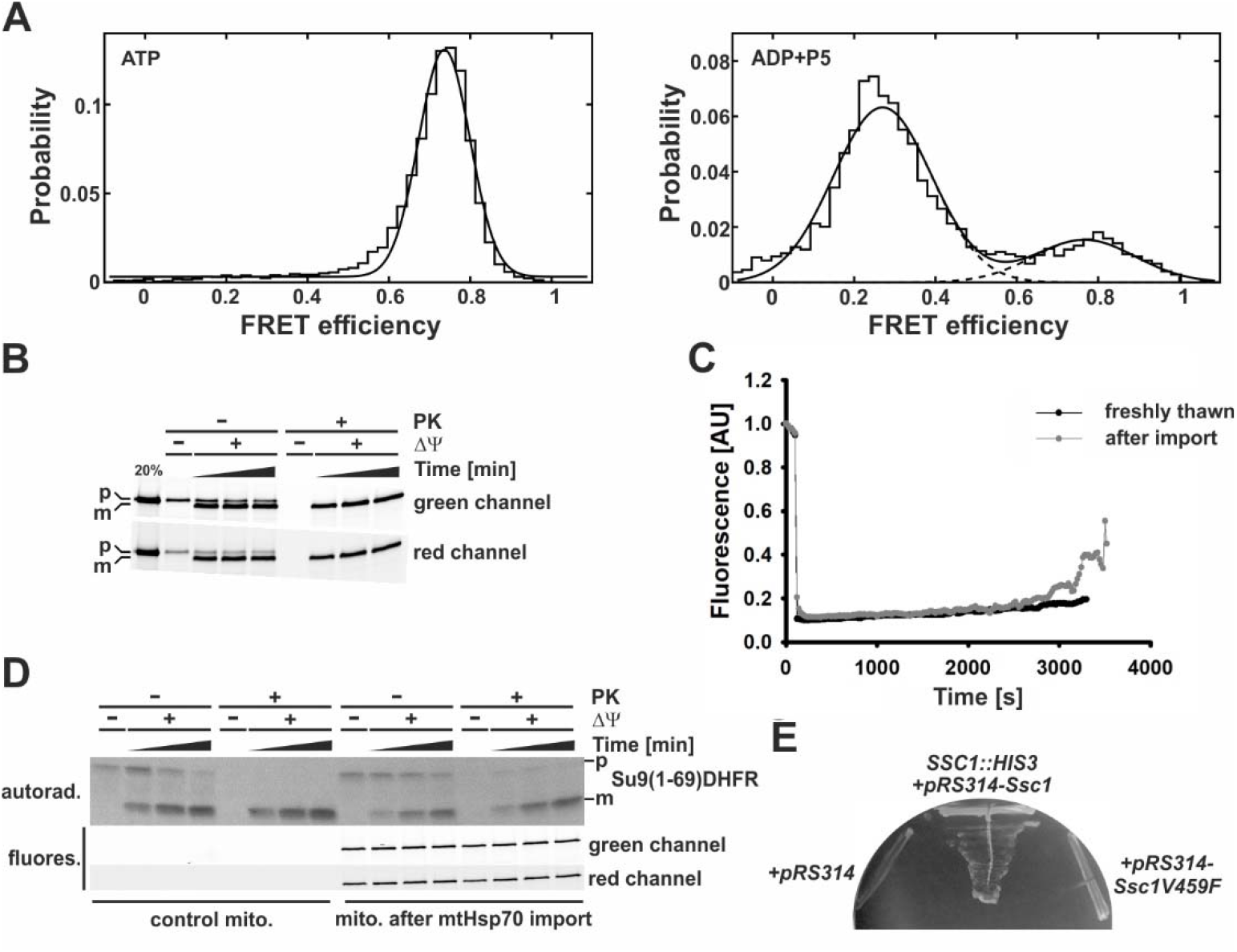
Control experiments for the smFRET analysis in intact mitochondria. (**A**) Single-molecule FRET histograms from solution-based *in vitro* burst measurements of mtHsp70 in the presence of ATP (left), and ADP and substrate P5 (right). Dashed lines indicate single Gaussians of the subpopulations of the Gaussian fit (solid line). (**B**) Uptake of labeled mtHsp70 into isolated mitochondria. The recombinantly expressed and purified precursor form of mtHsp70 was labelled with donor (Atto532) and acceptor (Atto647) fluorophores and incubated with isolated mitochondria in the presence of membrane potential (ΔΨ). After 5, 15 and 40min, aliquots were taken out and import was stopped. As a control, one sample was incubated with deenergized mitochondria. One half of the samples was treated with proteinase K to degrade nonimported material. Mitochondria were reisolated and samples analyzed using SDS-PAGE followed by detection of fluorescent signals in the red and green channels using a Typhoon FLA 9500. (**C**) The membrane potential (ΔΨ) of isolated mitochondria, after import of fluorescently labeled mtHsp70 or freshly taken from - 80°C freezer, was measured by the fluorescence quenching of DiSC_3_(5). A drop in the ΔΨ of mitochondria (i.e. an increase in fluorescence) measured during import with fluorescently labelled mtHsp70 started after ca. 45 min. (**D**) [^35^S]-labeled precursor protein Su9(1-69)DHFR was incubated with isolated mitochondria, after import of fluorescently labeled mtHsp70 or mitochondria freshly taken from the -80°C freezer, in the presence of ATP, NADH and an ATP-regenerating system. After 1, 3 and 9min, aliquots were taken out and import was stopped. In one sample, the membrane potential was dissipated prior to addition of the precursor protein. One half of the samples were treated with proteinase K (PK) to degrade nonimported material. Mitochondria were reisolated and samples analyzed using SDS-PAGE followed by detection of fluorescent signals using a Typhoon FLA 9500 and radioactive signals by autoradiography. p, precursor and m, mature forms of Su9(1-69)DHFR. (**E**) Haploid yeast strain with chromosomal deletion of *SSC1*, encoding mtHsp70, and rescued with the wild-type copy of mtHsp70 encoded on a *URA* plasmid was transformed with centromeric yeast plasmids pRS314 encoding either wt or the V459F mutant version of mtHsp70 under the control of its endogenous promoter and 3’
sUTR. An empty plasmid was used as a negative control. Cells were plated on medium containing 5-fluoroorotic acid, which selects against the *URA* plasmid and thereby enables one to assess the functionality of mtHsp70 encoded by the pRS314 plasmids.

## Materials and Methods

### Yeast strains and growth conditions

A list of yeast strains used in this study is given in Table S1. Haploid yeast strain YPH499 was used as wild type (*33*). For isolation of mitochondria, cells were grown in lactate medium containing 0.1% glucose at 30°C.

Yeast strain expressing matrix targeted DHFR_mut_ from a *GAL* promoter and the corresponding control strain carrying an empty pYES2 plasmid were described previously(*20*). A list of plasmids used in this study is given in Table S2. Cells were grown in selective lactate medium containing 0.1% glucose at 30°C and diluted twice a day, once in the morning and once in the evening, so that they always remain in the logarithmic growth phase. To induce expression of mDHFR_mut_, 0.5% galactose was added to the growing cultures. Mitochondria were isolated from cells two hours after addition of galactose. The Ssc1 (major mtHsp70 in yeast) shuffling strain was made by transforming YPH499 with a *URA3* plasmid pVT-U (*34*) encoding the wild-type Ssc1 and subsequently deleting the chromosomal copy of *SSC1* with a *HIS3* marker using homologous recombination. To analyze the ability of the Ssc1V459F mutant to support growth of yeast cells, the Ssc1 shuffling strain was transformed with the yeast centromeric plasmid pRS314 (*33*) encoding Ssc1V459F under the control of the endogenous Ssc1 promoter and 3’
sUTR. Empty plasmid and plasmid encoding the wild-type Ssc1 were used as negative and positive controls, respectively. The ability of Ssc1 variants encoded on the pRS314 plasmids to support growth of yeast cells was analyzed on plates containing 5-fluoroorotic acid, which selects against the *URA* plasmid.

### Cloning, expression, purification and labelling of recombinant proteins

pET-Duet plasmids for recombinant expression of the mature Ssc1 (residues 24 to 645) and its two-cysteines variant (D341C, I448C) along with the His-tagged Hep1 were described before (*26, 30*). Purification and labelling of mature Ssc1 was done as described before (*26*) with the exception that ATTO 647 (ATTO-TECH GmbH, Siegen, Germany), instead of ATTO 647N, was used as the acceptor fluorophore.

For expression of the precursor form of Ssc1 (residues1 to 645), the C-terminally His_6_-tagged version of Ssc1(D341C,I448C) was cloned into pET-Duet plasmid between NdeI and XhoI restriction sites. The primers used for PCR amplification are given in Table S3 and the pRS314 plasmid encoding the two-cysteine variant of Ssc1(D341C, I448C) under the control of the endogenous *SSC1* promoter and 3’
sUTR was used as the template. The thus obtained pET-Duet plasmid served as the template to generate the V459F variant by site directed mutagenesis using the primers described in Table S3. Plasmids were transformed into electrocompetent BL21(DE3) *ΔdnaK::52* cells (a kind gift of Dr. Meyer, ZMBH, Heidelberg, Germany) and the cells grown in LB medium supplemented with 100 µg/mL ampicillin overnight at 37°C. Overnight cultures were diluted into fresh LB-Amp medium and grown at 37°C till they reached OD_600_ of ca. 0.5. The expression of proteins was induced by the addition of 0.5mM IPTG and the cells were grown for an additional 3h at 37°C. Cells were harvested by centrifugation (3000g, 10min, 4°C) and kept at -20°C until use. The precursor form of Ssc1 was insoluble after recombinant expression, irrespective of whether Hep1 was co-expressed or not, and was therefore purified from inclusion bodies. To this end, the cell pellet was resuspended in 50 mM Na-phosphate buffer at pH 7.5, 150 mM NaCl, 2 mM DTT, 1 mM PMSF and the cells lysed by sonication (15 cycles of 12s sonication, 18s break; 80% duty cycle, setting 5; Branson sonifier). Soluble proteins were separated by centrifugation (27000g, 15 min, 4°C). Pellet-containing Ssc1 was resuspended in TRIS/urea buffer (50 mM TRIS/HCl, 3 M urea, 2 mM DTT, 20 mM imidazole, pH 8.0) and incubated on a rotating platform for 1h at 25°C. The sample was centrifuged (27000g, 15 min, RT) and the supernatant, containing Ssc1, was passed through a 1 mL NiNTA-agarose column (Qiagen) preequilibrated with the TRIS/urea buffer. The column was washed with 30 column volumes of the TRIS/urea buffer before Ssc1 was eluted using TRIS/urea buffer containing 300 mM imidazole. Imidazole was removed from the sample by exchanging the buffer on a PD10 column (Cytiva) equilibrated with 50 mM TRIS/HCl, 3 M urea and 2 mM DTT at pH 8.0. Protein was concentrated to ca. 3 mg/mL (∼40 µM) using Amicon 10K concentrators (Merck) and stored at -80°C till use.

For labeling, purified proteins were taken from -80°C storage and DTT removed from the samples by buffer exchange on a PD10 MiniTrap G25 column (Cytiva) equilibrated with labeling buffer (50 mM TRIS/HCl, 3 M urea, pH 7.0). The proteins were immediately labeled with ATTO 532 maleimide and ATTO 647 maleimide (both ATTO-TECH GmbH) as donor and acceptor fluorophores, respectively. For this, the protein, and donor and acceptor dyes were mixed in a 1:2:5 molar ratio and incubated on an overhead rotator for 3h at 4°C. Free dyes were removed on a PD10 MiniTrap G25 column (Cytiva) equilibrated with labeling buffer and the labeled proteins stored in aliquots at -80°C.

### Sample preparation for in organello smFRET measurements

For import of the fluorescently labelled precursor form of Ssc1, isolated wild-type mitochondria (200 µg) were resuspended in 400 µL SI buffer (50 mM HEPES/KOH, 0.6 M sorbitol, 75 mM KCl, 10 mM Mg-acetate, 2 mM KH_2_PO_4_, 2.5 mM EDTA, 2.5 mM MnCl_2_, pH 7.2, 0.5 mg/mL fatty acid-free BSA) and incubated at 25°C for 10 min. ATP (2.5 mM), NADH (3.3 mM), creatine-phosphate (10 mM) and creatine kinase (0.1 mg/mL) were then added and incubation continued for another 3 min. The fluorescently-labelled precursor form of Ssc1 was then added to energized mitochondria and imported for 10 min. For every new batch of mitochondria and labelled protein, different dilutions of labelled protein were tested so that, on average, less than one molecule of protein is imported per isolated mitochondrion. To stop further import, the import mix was diluted with 1 mL ice-cold SH buffer (0.6 M sorbitol, 20 mM HEPES/KOH, pH 7.2) and proteinase K (0.1 mg/mL) was added to degrade nonimported material. After 5 min on ice, PMSF (2 mM) was added to stop the digestion and the sample incubated for a further 5 min on ice. Mitochondria were reisolated (13600g, 10 min, 4°C), resuspended in 1 mL SH buffer containing 3 mg/mL fatty acid-free BSA, reisolated again and finally resuspended in 20 µL SH buffer and kept on ice.

Glass cover slides (Marienfeld, High Precision Microscope Cover Glasses, 24 × 60 mm, 170±5 µm, No. 1.5H) were cleaned with acetone (HPLC grade) and pure ethanol in a sonication bath. After drying, the surface was coated with 5-10 v% PEG-Silane (ABCR, 3-[Methoxy(polyethyleneoxy)propyl]trimethoxysilane, 90%, 6-9 PE-units) and 3-5 mg Silane-PEG-Biotin (Nanocs, 3400 Da) in a solution of 50 mL toluene at 55 °C. A piece of red silicon sheet (CoverWell Press-to-seal, 2.4 mm thickness) with a hole (Ø 8 mm) in the middle was put on the washed and dried coverslip and fixed with a coat of epoxy glue (TOOLCRAFT 5 min Epoxy) around the edges. The hole acts as the sample chamber.

The sample chamber was incubated with 100 µL of 0.2 mg/mL streptavidin (Sigma-Aldrich, from *Streptomyces avidinii*) for 15 min. Afterwards, the chamber was washed three times with PBS. In the next step, biotinylated, affinity-purified antibody against the N-terminal peptide of Tom22 was added to the chamber. After 15 min incubation, the chamber was washed three times with PBS and once with SI buffer. The prepared batch of mitochondria with imported fluorescently labelled Ssc1 was taken off the ice, mixed with 80 µL SI buffer at room temperature (∼25 °C) and added to the chamber. The two experiments, import of fluorescently labelled Ssc1 and activation of cover slides with streptavidin/biotinylated antibodies, were performed such that both samples were ready at the same time. After 10 min incubation at room temperature, the chamber was washed three times with SI buffer before 100 µL SI buffer containing 8 mM ATP, 8 mM NADH, 20 mM creatine-phosphate and 0.2 mg/mL creatine kinase were added. The sample was measured immediately for approximately 25 min. Under these experimental conditions, the mitochondria remain energized for about 45 min at room temperature (fig. S2C). Hence, the sample was discarded after 40 min had passed since removal of the mitochondria from the ice.

### HILO measurements

Objective-type single-molecule Förster Resonance Energy Transfer (smFRET) experiments were carried out on a home-built three-color single-molecule total internal reflection /widefield fluorescence microscope using highly inclined and laminated optical sheet (HILO) illumination with alternating laser excitation (ALEX) (*35-37*). The sample was illuminated with blue laser light (491 nm, Cobolat Calypso, 50mW) at 18 mW for the initial 50 frames to allow localization of the mitochondria by detection of their autofluorescence. Subsequently, the sample was illuminated with alternating green (532 nm, Cobolt Samba, 100 mW) and red (647 nm, Cobolt MLD, 120 mW) excitation using 30 mW and 20 mW measured at the sample, respectively, for 600 frames each, leading to a total of 1250 frames. The exposure time was 25 ms or 50 ms per frame, depending on the measurement. An overlap of mitochondria and both fluorophores indicates double-labeled protein inside a mitochondrion. From these measurements, single-molecule time traces were extracted. The acquired movies were analyzed using a home-written Matlab based software. The data were corrected with globally determined correction factors (the detection correction factor (gamma) = 0.97536; spectral crosstalk (beta) = 0.10802).

### PIE-MFD measurements

*In vitro* smFRET measurements of freely diffusing Ssc1 were performed on a dual-color confocal microscope with pulsed-interleaved excitation (*38, 39*) and multi-parameter fluorescence detection (*40, 41*). The power of the two lasers having wavelengths of 532 nm (PicoTA Toptica) and 640 nm (PicoQuant LDH-D-C-640) was set to 100 µW measured before the objective. The acquired data was analyzed with PAM, a Matlab-based software (*42*). For the *in vitro* smFRET measurements, labelled mature Ssc1 was diluted to 10-50 pM concentration in measurement buffer (20 mM Tris-HCl pH 7.5, 80 mM KCl, 5 mM MgCl_2_) such that only one protein diffuses through the confocal volume at a time. The goal was to have 2-5 bursts per second. Samples contained either 1 mM ATP or 1 mM ADP and 50 µM model substrate peptide P5 (CALLSAPRR).

### Solution measurements after lysis of mitochondria

The mitochondria with imported labelled Ssc1 were taken from the ice and added to a probe chamber as for TIRF measurements. After washing with SI buffer, measurement buffer (20 mM Tris-HCl pH 7.5, 80 mM KCl, 5 mM MgCl_2_) containing 0.5% Triton X-100, 5 mM ATP and 1 mM PMSF was added to the mitochondria. After 3 min, the solution was taken out of the probe chambers and diluted to single-molecule concentrations in measurement buffer (20 mM Tris-HCl pH 7.5, 80 mM KCl, 5 mM MgCl_2_) with 5 mM ATP.

### Fluorescence microscopy

For imaging, yeast cells were grown to an early logarithmic phase in synthetic lactate medium lacking uracil and supplemented with 0.1% (w/v) glucose and 10 mg/L adenine at 30°C. To induce the expression of unfolded proteins, 0.5% (w/v) galactose was added to the growing cultures. After two hours, cells were collected and stained with 50 nM Mitotracker Red CMXRos (Life Technologies) for 30 min and immobilized on agar pads for imaging.

Microscopy was performed at 30°C on a Nikon Ti2-Eclipse microscope. All images were postprocessed by deconvolution using the NIS-Elements software (Nikon) and analyzed using Fiji (*43*). A blind quantification was performed by grouping cells according to their mitochondrial morphology as tubular, aggregated or fragmented. About 400 cells per strain were analyzed.

### Coimmunoprecipitation

Mitochondria (1 mg/mL) were solubilized with 1% digitonin in solubilization buffer (20 mM HEPES/KOH, 150 mM NaCl, 10% glycerol (v/v), 0.1 mM EDTA, 2 mM PMSF, pH 7.4) on an overhead roller for 15 min at 4°C. After a clarifying spin (124500g, 20 min, 2°C), solubilized mitochondria were added to ProteinA-Sepharose CL4B beads (Cytiva) with prebound affinity purified antibodies against Ssc1 or antibodies from a preimmune serum, as a negative control. Samples were incubated on an overhead roller for 45 min at 4°C. After three washing steps with the solubilization buffer containing 0.05% digitonin, specifically bound proteins were eluted with reducing Laemmli buffer for 3 min at 95°C. Samples were analyzed by SDS-PAGE followed by immunoblotting. A list of antibodies used in this study is given in Table S4.

Six independent coimmunoprecipitation experiments performed with two different batches of isolated mitochondria were quantified. The signal intensities were corrected for the background and normalized to the amount of mtHsp70 precipitated from the same mitochondria. The amount of immunoprecipitated material in control mitochondria was set to 100%. Image analysis was done using Fiji and calculations were done using Excel and RStudio.

### Miscellaneous

The following procedures were performed using previously published protocols: preparation of yeast total cell extracts (*44*), isolation of mitochondria (*45*), synthesis of radiolabelled precursor proteins (*45*), protein import into mitochondria (*45*) and measurement of membrane potential (*46*). Chemical crosslinking was done as previously described (*31*) with the exception that the final concentration of the crosslinker was 150 µM, instead of 75 µM.

---

**Table S1.** *Saccharomyces cerevisiae* strains used in this study.

| Strain | Genetic background | Source |
| --- | --- | --- |
| YPH499 (WT) | *MATa ura3-52 lys2-801<sup>amber</sup> ade2-101<sup>ochre</sup> trp1-*<br>*Δ63 his3-Δ200 leu2-Δ1* | (33) |
| WT+pYES2 | YPH499 pYES2 | (19) |
| WT+pYES2-<br>mDHFR<sub>mut</sub> | YPH499 pYES2-mDHFR<sub>mut</sub> | (19) |
| Ssc1 shuffling strain | YPH499 *Δssc1::HIS3* pVTU-Ssc1 | This study |

**Table S2.** Plasmids used in this study.

| Plasmid | Source |
| --- | --- |
| pETDuet-Ssc1(24-645)(D341C,I448C)+HisHep1 | (26) |
| pETDuet-Ssc1(1-645)(D341C,I448C)His<sub>6</sub> | This study |
| pETDuet-Ssc1(1-645)(D341C,I448C,V459F)His<sub>6</sub> | This study |
| pVT-U | (34) |
| pVT-U-Ssc1 | This study |
| pFA6a-His3MX6 | (47) |
| pRS314 | (33) |
| pRS314-pSsc1f | This study |
| pRS314-pSsc1V459Ff | This study |

**Table S3.**
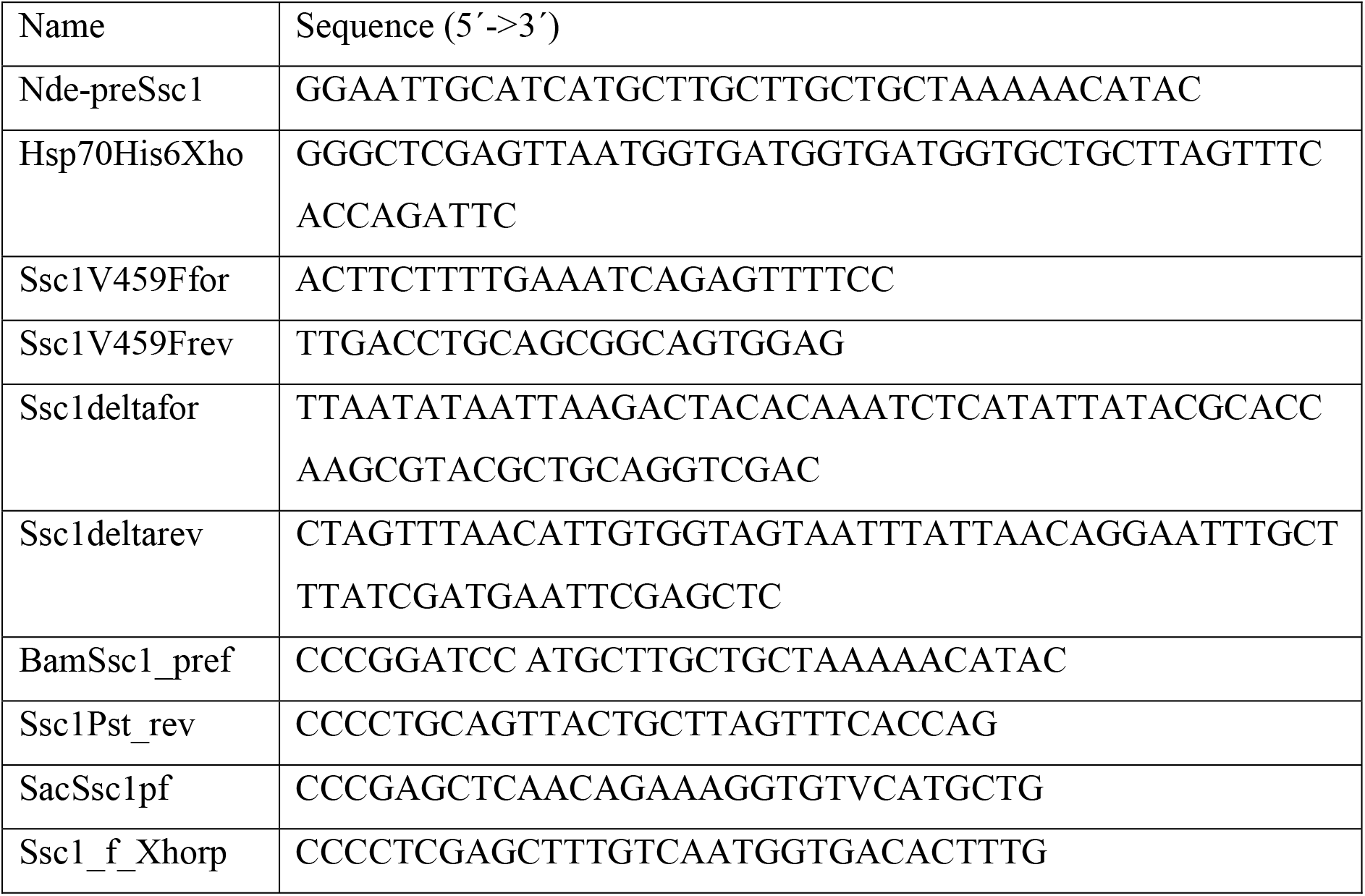
Primers used in this study.

**Table S4.** Antibodies used in this study.

| Antigen | Source | No. |
| --- | --- | --- |
| DHFR | Invitrogen | MA5-24945 |
| Hsp60 | Neupert Lab antibodies | 80322 |
| mtHsp70 | Neupert Lab antibodies | 347 (affinity purified) |
| Mdj1 | This study | 548 |
| Mge1 | Neupert Lab antibodies | 366 |
| Hep1 | (30) | 316 |
| Ssq1 | (48) | 181 |
| Yme1 | (19) | Yme1Cpep (affinity purified) |
| Phb2 | Neupert Lab antibodies | 429 |
| Tim44 | (49) | 388 (affinity purified) |
| Tim14 | (31) | 313 (affinity purified) |
| Tim17 | Neupert Lab antibodies | Tim17Cpep (affinity purified) |
| Tim23 | Neupert Lab antibodies | Tim23Npep (affinity purified) |
| Porin | Neupert Lab antibodies | 87118 |
| Tom22 | Neupert Lab antibodies | Tom22Npep (affinity purified) |
| Tom40 | This study | 547 (affinity purified) |
| Cytochrome c | Neupert Lab antibodies | 442 |
| Cox2 | Neupert Lab antibodies | 46 |
| F1β | Neupert Lab antibodies | 421 |
| Aconitase | Neupert Lab antibodies | 206 |

